# Human Fcγ-receptors selectively respond to C-reactive protein (CRP) isoforms

**DOI:** 10.1101/2025.03.23.644781

**Authors:** Anna Henning, Johanna Seer, Johannes Zeller, Karlheinz Peter, Julia Thomé, Philipp Kolb, Steffen U Eisenhardt, Katja Hoffmann, Hartmut Hengel

## Abstract

The pentameric C-reactive protein (pCRP), an acute-phase protein, binds to lysophosphatidylcholine (LPC) displayed on the surface of dying cells and microorganisms to activate the complement system and to opsonize immune cells via Fcγ-receptors (FcγRs). Members of the FcγR family are characterized by the recognition of the Fc part of IgG antibodies. We utilized a mouse thymoma BW5147 reporter cell panel stably expressing chimeric human FcγR-CD3ζ-chain receptors to define the molecular requirements for FcγR crosslinking by C-reactive protein (CRP). Applying this approach, we show a robust activation of CD64/FcγRI and CD32a/FcγRIIa by immobilized CRP isoforms as well as triggering of inhibitory CD32b/FcγRIIb. Of note, activation of FcγRIIa was restricted to the 131R allelic variant but not observed with 131H. In contrast, FcγRIII isoforms CD16aF, CD16aV and CD16b were not activated by pCRP, although binding of CRP isoforms to FcγRIII was detectable. Activation of FcγRs by free pCRP in solution phase was considerably lower than with immobilized pCRP on hydrophilic plastic surfaces and readily abolished by IgG at serum level concentrations, whereas it was enhanced by the addition of streptococci. The types of FcγRs mainly responding to pCRP in solution phase (CD64/FcγRI and CD32aR/FcγRIIaR) clearly differed from FcγRs responding to soluble multimeric IgG complexes (i.e., CD16aV/FcγRIIIaV and CD32aH/FcγRIIaH). Compared to pCRP, monomeric CRP (mCRP) showed lower levels of activation in those selective FcγRs. FcγR activation was linked to recognition by conformation-dependent CRP antibodies. Unmasking of the mAb 9C9-defined neoepitope in pCRP* correlated with the triggering of FcγRs, indicating that pCRP* is the major FcγR-activating CRP conformation. The assay provides a novel, scalable approach to determine the molecular properties of CRP as a physiological ligand of FcγR-mediated bioactivities.

**Scope statement:** Fcγ receptors (FcγRs) are important immune regulators that come in different variants and combinations, making it difficult to predict which components will ultimately lead to immunological effector functions. Classical FcγRs are defined by their recognition of IgG-Fc, while other ligands, such as C-reactive protein (CRP), are often neglected. Circulating concentrations of CRP, an acute phase protein, are elevated during inflammatory responses. As a pattern recognition receptor, CRP binds to lysophosphatidylcholine expressed on the surface of dying cells and microbes in order to activate the complement system via C1q.

We have established a reporter cell assay platform that goes beyond ligand binding and takes a deeper look at the activation outcome(s) by CRP compared with IgG-Fc. This is the first comprehensive study defining CRP-responsive vs non-responsive FcγRs and investigating the interaction of FcγRs with CRP isoforms (pCRP/pCRP*/mCRP). We distinguish binding from receptor triggering using reporter cells stably expressing a chimeric FcγR-CD3ζ chain, thereby defining the molecular requirements for FcγR cross-linking by CRP. The assay provides a novel, sensitive and scalable approach to the properties of CRP as a ligand inducing FcγR-mediated bioactivities.

## 1 Introduction

C-reactive protein (CRP) is a pattern recognition molecule and prototypical acute-phase protein. It is widely used as a marker of acute inflammation in patients. CRP is a member of the pentraxin family and synthesized mainly by hepatocytes (1–3). The secreted CRP molecule consists of five identical non-covalently linked non-glycosylated protomers of ∼23 kDa each. These protomers are aligned in planar symmetry to form a donut-shaped ring (4). This ring comprises two faces, i.e., the complement C1q or Fcγ-receptor (FcγR) binding ‘effector’ A-face and the ligand binding B-face. Phosphocholine (PC) head groups expressed on bacterial cell walls and damaged host cell membranes (5,6) are the prototypic ligands for CRP. PC is bound in a calcium-dependent manner via the phosphocholine binding pockets expressed on the B-face. The opposite A-face of the pentamer contains overlapping binding sites for C1q and FcγRs, so that the two interaction domains are considered to be mutually exclusive (7).

Traditionally, a distinction is made between at least two main conformational isoforms of CRP: The circulating native, pentameric CRP (pCRP) and the monomeric isoform (mCRP), which is ultimately formed by dissociation of the pentameric molecule. Under experimental conditions, this process can be initiated by exposure to heat, acid or urea and leads to the exposure of neoepitopes on the CRP molecule that are inaccessible in the native pentameric form (8–10). *In vivo*, the dissociation process is observed on PC-rich membranes of activated platelets, monocytes or endothelial cells, by interaction with misfolded proteins and by mechanical stress in stenosed vessels (10–14). In contrast to pCRP, mCRP is insoluble and considered a pro-inflammatory, tissue- or cell-bound isoform of CRP found deposited to local inflammation. A third intermediate isoform of CRP, pCRP*, has only recently been described (10). Binding of pCRP to microparticles containing PC head groups released by activated cells leads to a conformational change in the structure of pCRP: the neoepitopes responsible for C1q and FcγR binding that are accessible in mCRP are also exposed in pCRP*, but unlike mCRP, the overall pentameric symmetry is preserved. Exposure of the neoepitopes facilitates C1q binding and complement activation, with the result that pCRP* can increase tissue inflammation (10).

FcRs form a vital link between humoral and cellular immunity: They recognize the Fc region of antibodies bound to antigens via their Fab region. IgG-binding FcγRs belong to the immunoglobulin superfamily expressed on most immune effector cells. They can be divided into activating and inhibitory FcγRs. Both FcγR types are often expressed on the same cell and form a binary system integrating activating and inhibitory signals (15). FcγRI (CD64), FcγRIIa (CD32aH/R), FcγRIIc (CD32c), FcγRIIIa (CD16aF/V) and FcγRIIIb (CD16b) are activating FcγRs and (except for FcγRIIIb) signal via immunoreceptor tyrosine based activating motifs (ITAMs) in their cytoplasmic regions (16,17). FcγRIIb (CD32b), the only inhibitory FcγR, signals via an immunoreceptor tyrosine based inhibitory motif (ITIM) (18). FcγRI (CD64) is the high affinity receptor for IgG, whereas all other FcγRs have low to medium affinity to monomeric IgG (19). Binding of either immobilized or multimeric soluble immune complexes (ICs) to FcγRs leads to various effector functions that depend on the FcγRs expressed and the type of immune effector cell affected and include antibody-dependent cellular cytotoxicity (ADCC), antibody-dependent cellular phagocytosis (ADCP), cytokine release, oxidative burst and apoptosis (18,20).

Recognition of pCRP by FcγRI (CD64) and FcγRIIa (CD32a) was first demonstrated by flow cytometry using transfected COS-cells and monoclonal antibodies (21,22). Later studies characterized the binding of pCRP to FcγRs in antibody- and label-free setups. FcγRIIa was found to dock diagonally to two of the five pentraxin subunits on the effector face with its D1 and D2 domains, ensuring a one-to-one binding stoichiometry with no significant conformational changes (23). Binding of pCRP was observed not only with FcγRI (CD64) and FcγRIIa/b (CD32), but also with FcγRIII (CD16) (23,24). Binding affinities of pCRP to FcγRs are in a similar range (24) and comparable to IgG binding to low affinity FcγRs (25). Pentraxin binding sites partially overlap with IgG binding sites on FcγRs, suggesting competitive binding (23). Binding of pCRP to FcγRs leads to opsonization, cytokine production and enhancement of phagocytosis (23,26,27).

Many aspects of CRP-FcγR interaction remain controversial. Preferential binding of pCRP to the 131R allelic variant of FcγRIIa compared to 131H has been considered certain for decades and various clinical observations have been attributed to this difference (28,29). However, recent contrary observations of a potential difference in pCRP binding to FcγRIIa-H/R131 have been made in antibody free setups (23,24). Whilst several studies investigate the interaction of different conformational isoforms of CRP (pCRP/pCRP*/mCRP) and C1q, little is known regarding the impact of the CRP isoforms on FcγR activation (10,11,12,30,31). Neither a clearly pro-nor anti-inflammatory role can be attributed to the CRP–FcγR interaction, as both pro- and anti-inflammatory cytokine expression have been reported (23,27,32). The precise contribution of CRP to immune complex-mediated diseases and the intricate interplay between CRP, IgG and FcγRs remains to be elucidated (24,33–35).

The BW5147-FcγRζ reporter assay panel is based on mouse BW5147 thymoma cells stably transduced with the extracellular domain of individual human, rhesus, or mouse FcγRs (e.g., human FcyRI/IIaH/IIaR/IIb/IIIaF/IIIaV/IIIb), allowing for convenient, quantifiable, and high-throughput analysis of FcγR activation by IgG (33–35). The assay has been established for immobilized IgG, multimeric immune complexes in solution phase (sICs) and recombinant Fc-fusion therapeutics mediating activation of FcγRs (20,33,36,37). Unlike the variety of FcγRs found on primary immune cells, each setup contains only one FcγR, allowing clear attribution of the observed activation. FcγR ectodomains are coupled to the signaling CD3ζ chain of the TCR, leading to mouse IL-2 (mIL-2) production upon receptor cross-linking and activation of the reporter cell. Here, we modified the test system to detect activation mediated by distinct human CRP isoforms and to compare CRP-dependent with IgG-mediated activation. While binding of pCRP has been investigated for individual FcγRs, pCRP-dependent activation has solely been examined in complex settings with several FcγRs and/or more than one cell type present. In this study, the reductionistic setup of the BW5147-FcγRζ reporter assay allowed for comparing specific FcγR binding to distinct CRP isoforms with subsequent FcγR crosslinking and activation, as well as interactions of CRP with IgG and soluble immune complexes which are independent ligands of FcγRs. The BW5147-FcγRζ test system distinguished CRP-responsive (CD64/FcγRI, CD32aR/FcγRIIaR, and CD32b/FcγRIIb) from non-responsive human FcγRs and revealed a clear allele-dependent activation pattern of CD32a/FcγRIIa by CRP (131R>>H). Triggering of FcγRs was achieved by either soluble or immobilized pCRP or mCRP ligand, with immobilized pCRP showing highest triggering efficacy. Interestingly, effective pCRP signaling via FcγRs was associated with conformational unmasking of the pCRP*/mCRP neoepitope as detected by mAb clone 9C9 and activation caused by pCRP was stronger than for mCRP, suggesting pCRP* as the major FcγR activator (10).

## 2 Materials and Methods

### 2.1 CRP preparation and detection, IgG source and sICs preparation

Highly purified human CRP from pleural fluid/ascites and recombinant CRP produced in E. coli (C7907-26 and C7907-03C) was purchased from US Biological Life Sciences (Salem, Massachusetts, USA) mCRP was prepared from purified pCRP as described previously (38) and concentrations of pCRP and mCRP were measured using Qubit Fluorometric Quantitation (Thermo Fisher Scientific, Waltham, MA, USA). Streptococcus pneumoniae serotype 27 was kindly provided by Dr. Mark van der Linden, Head of the National Reference Center for streptococci, Department of Medical Microbiology, University Hospital (RWTH, Aachen, Germany). To form CRP-streptococci complexes, 10 µl of suspended streptococci were added to 20/10/5 µg of CRP (39,40).

Synthetic sICs formed by 25 nM Infliximab (149.1 kDa) and 50 nM TNFα monomer (17.5 kDa) to ensure a 1:1 stoichiometry were produced as described previously (20). sICs and CRP-streptococci complexes were incubated for two hours at room temperature (RT) prior to being used in the experiment. Polyclonal goat anti-human CRP antibody (A80-125A) was purchased from Bethyl (Montgomery, Texas, USA), monoclonal conformation-specific antibodies binding pCRP and pCRP*/mCRP (clone 8D8 and 9C9, respectively) were kindly provided by Prof. Lawrence A. Potempa, College of Pharmacy, Roosevelt University, Schaumburg, IL, USA. LPS (LPS EB Standard, 5 mg, #tlrl-eblps, LPS *E. coli* O111:B4) was purchased from InvivoGen (San Diego, California, USA). Purified human IgG (cytotect®, Biotest, Dreieich, Germany), recombinant Rituximab IgG1 (humanized monoclonal; Roche, University Hospital Freiburg Pharmacy), and concentrated IgG1 (human IgG1 kappa, #I5154-1MG; Sigma-Aldrich, St. Louis, Missouri, USA) served as sources of IgG.

#### 2.1.1 BW5147 cell culture

The murine T lymphoblast cell line BW5147 (TIB-47™; ATTC, Manassas, VA, USA) was maintained in RPMI 1640 medium (“RPMI BW medium”, GlutaMAX™; Gibco Life Technologies, Carlsbad, California, USA) supplemented with 10% (v/v) heat-inactivated FCS (Biochrom, Berlin, Germany), 1% (v/v) Pen-Strep (Gibco Life Technologies), 1% (v/v) sodium pyruvate (100 mM, Gibco Life Technologies), and 0.1% (v/v) β-mercaptoethanol (Sigma-Aldrich). Cells were cultured at 37°C with 5% CO₂ and split based on their growth rate. Cells were maintained at a density of 2×10⁵/ml to 1×10⁶/ml. For the FcγR activation assay, cells were seeded at 2–3 × 10⁵ cells/mL one day prior to the experiment, resulting in a density of 4–6 × 10⁵ cells/mL at the time of the assay. Cells were tested regularly for mycoplasma contamination using PCR (sense (#1427): 5’-GGGAGCAAACAGGATTAGATACCCT-3’; antisense (#1428): 5’-TGCACCATCTGTCACTCTGTTAACCTC-3’) with Kapa Polymerase (Peqlab, Erlangen, Germany #KK3604).

### 2.2 Flow Cytometry

BW5147 cells (100,000) were counted using a Countess® II automated cell counter and centrifuged at 1,000 rpm at RT for six minutes. Cells were washed twice in 100 µl FACS buffer (PBS (Dulbecco’s PBS, Gibco Life Technologies) with 3% (v/v) heat-inactivated FCS (Sigma-Aldrich)) on ice and centrifuged at 1,400 rpm at 4°C for five minutes. Each sample (100,000 BW5147 cells in 300 µl) was incubated with v/v 1:100 mouse-anti-human-CD16 allophycocyanin (APC) (FcγRIII, clone B73.1), mouse-anti-human-CD32 APC (FcγRII, clone FUN2), mouse-anti-human-CD64 APC (FcγRI, clone S18012C), or mouse-anti-human-CD99 APC (MIC2, clone hec2)-APC (200 µg/ml, BioLegend, San Diego, California, USA; cat. #360705, #303207, #399509, #398203, respectively) antibodies on ice for one hour. Respective anti-FcγR antibodies on BWCD99 cells or unstained BW parental cells served as negative controls for background antibody binding. Cells were washed three times and transferred to FACS round-bottom polystyrene test tubes (Falcon®) containing 200 µl FACS buffer. Samples were kept on ice until analysis using a BD LSR Fortessa™ Cell Analyzer (BD biosciences, Franklin Lakes, New Yersey, USA). A total of 20,000 events were measured per sample. Results were analyzed using FlowJo software (FlowJo LLC, Ashland, OR, USA), with gating applied to the main population (FSC/SSC gating). APC-A fluorescence was compared using histograms normalized to mode.

### 2.3 BW5147-FcγRζ reporter assay

The BW5147-FcγRζ-cell reporter assay, i.e., mouse BW5147 hybridoma cells stably expressing chimeric FcγR-CD3ζ chain molecules consisting of an extracellular domain of human FcγRs fused to the transmembrane and intracellular domains of the mouse CD3ζ chain (32), enables analysis of IgG-mediated activation of individual subclasses of human FcγRs. The general procedure of the BW5147-FcγRζ reporter assay was utilized as described before (33) and modified to analyze human CRP-mediated activation (20,33,37). In brief, BW5147 cells that are stably transduced with the extracellular domain of one of the human FcγRs (FcγRI, IIaH, IIaR, IIb, IIIaF, IIIaV, IIIb) or with human CD99 as a negative control were used. The human FcγRζ-receptor ectodomain is fused to the signaling CD3ζ-chain of the mouse T cell receptor (TCR), subsequently inducing mouse IL-2 (mIL-2) expression upon receptor crosslinking. In this assay, mIL-2 production is directly proportional to FcγR activation. mIL-2 levels were measured using a sandwich ELISA as described in detail below. For this project, the assay was modified to measure human CRP-dependent and IgG-mediated activation by FcγRζ-receptor crosslinking and as a positive control, respectively. BW5147 reporter cells were stably transduced via lentiviral transduction as described previously (20,33,34,41). FcγR expression was ensured by puromycin selection and two consecutive cell-sorting steps by FACS. BW5147-FcγRζ reporter assays were performed in 96-well ELISA MaxiSorp plates (Thermo Fisher Scientific, Immuno Platte F96 Maxi Pinchbar). For the ‘standard’ crosslinking assay, MaxiSorp plates were coated with graded concentrations of either IgG1 (human IgG1 kappa, #I5154-1MG, Sigma-Aldrich) or CRP isoforms in 50 µl PBS for one hour at 37°C with 5% CO₂ or overnight at 4°C. The protocol for the ‘in solution’ BW5147-FcγRζ reporter assay was adapted for CRP from the protocol established for soluble immune complexes (sICs) in our laboratory (20). ELISA wells were blocked by adding 300 µl ELISA blocking buffer (PBS with 10% (v/v) heat-inactivated FCS (Biochrom)) and incubating overnight at 4°C. sICs and complexes formed with pCRP and streptococci were incubated for two hours at RT. Complexes were added to ELISA wells in 100 µl of RPMI BW medium, followed by the addition of 100,000 BW5147-FcγRζ cells in another 100 µl of medium.

### 2.4 Sandwich mIL-2 ELISA

The level of mIL-2 secreted upon activation of BW5147 reporter cells was measured in a sandwich mIL-2 ELISA. ELISA MaxiSorp plates (Thermo Fisher Scientific) were coated with 50 µl of rat anti-mouse-IL2 antibody (1:500; 0.5 mg/ml; BD Pharmingen, BD biosciences, clone A85-1, #554424) in PBS -/- and incubated overnight at 4°C. Plates were washed and blocked as described above. Supernatants from the BW5147-FcγRζ reporter assay were transferred to the mIL-2-ELISA plates. Supernatants were incubated for 4 hours at RT on the ELISA plate, and wells were subsequently washed five times. 50 µl of biotinylated rat anti-mouse-IL2 (1:500; 0.5 mg/ml; BD Pharmingen, clone A85-1, #554426) in ELISA blocking buffer were added and incubated for 90 minutes at RT. Plates were washed five times, and 50 µl of Streptavidin-Peroxidase (1:1000; 1 mg/ml, Jackson ImmunoResearch, Philadelphia, PA, USA, #016-030-084) in blocking buffer was added for 30 minutes at RT. Wells were washed five times, and 50 µl of ELISA TMB 1-Step™ Ultra substrate solution (Thermo Fisher Scientific) was added, followed by 50 µl of 1 M H₂SO₄ to stop the reaction. Absorbance was measured using a Tecan ELISA Reader Infinite® M Plex (Tecan, Männedorf, Switzerland) at a wavelength of 450 nm and a reference wavelength of 620 nm.

### 2.5 Binding ELISA

Binding of recombinant His-tagged FcγRs (Sino Biological, Bejing, China, recombinant, HEK293/ECD, C-terminal polyhistidine tag: 1038-H08H1/10374-H08H1/10374-H08H/10259-H08H/10256-H08H/10389-H08C) to immobilized pCRP (US Biological Life Sciences) or IgG (human IgG1 kappa, # I5154-1MG, Sigma-Aldrich) was investigated by ELISA. Ninety-six-well ELISA MaxiSorp plates (Thermo Fisher Scientific) were coated with either pCRP, mCRP, or IgG1 overnight at 4°C, washed (PBS with 0.05% (v/v) Tween 20), and blocked with 300 µl of ELISA blocking buffer for one hour at RT. Blocking buffer was removed, and His-tagged FcγRs were added in 50 µl PBS. Binding was allowed to proceed overnight at 4°C. Subsequently, wells were washed five times, and 100 µl of blocking buffer/rabbit anti-His-antibody (1:5,000; 1 mg/ml, Bethyl: A190-114A) was added for overnight incubation at 4°C. Wells were washed five times, and goat anti-rabbit-peroxidase (POD) conjugated antibody (1:3,000; 1 mg/ml; Sigma-Aldrich; A0545) was added in 50 µl ELISA blocking buffer for one hour at 37°C. The ELISA readout using a Tecan ELISA Reader Infinite® M Plex at a wavelength of 450 nm and a reference wavelength of 620 nm was performed as described above. Binding assays in the ‘reverse’ setup were conducted following the same general procedure as described above. However, for this assay His-tagged hFcγRs were coated to ELISA wells in 50 µl PBS. Following the same blocking and washing steps as described above, IgG1, pCRP, or mCRP were added in 50 µl ELISA blocking buffer, and binding was detected using goat anti-hCRP antibody (1:3,000; 1 mg/ml; Bethyl: A80-125A) and donkey anti-goat (DAG) POD-conjugated antibody (DAG-POD; 1:5,000; 2,5mg/ml; Invitrogen, Waltham, Massachusetts, USA: A16005) for CRP (pCRP and mCRP) and goat anti-human-IgG-POD (1:3,000, 1 mg/ml; Rockland Immunochemical, Philadelphia, Pennsylvania, USA,, #109-035-003) for IgG1.

### 2.6 Semi-native PAGE and Coomassie

The structural integrity of pCRP as well as the monomeric form of mCRP was verified by semi-native gel electrophoresis as described previously (14). mCRP was generated by treating pCRP with 8 M urea in the presence of 10 mM EDTA for 2 hours at 37°C. mCRP was thoroughly dialyzed in low-salt phosphate buffer (10 mM Na₂HPO₄, 10 mM NaH₂PO₄, and 15 mM NaCl, pH 7.4). To confirm the use of pCRP or mCRP in the following assay setups, a pseudo-native SDS-PAGE and subsequent Coomassie staining or Western blot analysis with confirmation-specific CRP mAb was applied. In brief, samples were mixed with 15 µl of 1x sample buffer (1/20 of SDS as described in Lämmli-buffer, no DTT, no ß-ME), and pCRP or mCRP as indicated (10 µg, 5 µg or 3 µg). Samples were left without heating or boiling and loaded onto 10% PAA-Gel (all gel components and 1-Lämmli running buffer only with 1/20 of 20% SDS; final SDS concentration 1%). The gel was either directly stained with Coomassie brilliant blue solution and destained with water, or transferred onto a nitrocellulose membrane for Western blot analysis.

### 2.6 Statistical analyses and Graph modeling

Statistical analyses were performed using GraphPad Prism software (v9) and appropriate tests (Standard deviation; Ordinary One-way ANOVA for univariate comparison, and Two-way ANOVA for multivariate comparison followed by Tukey’s or Dunnett’s multiple comparisons test to assess significance; Area under the curve with standard error to compare activation patterns for multiple concentrations in titration setups). Generally, a significance level of p < 0.05 was applied. Higher p-values were considered not significant (ns) and are indicated as such on the graph, whereas p-values are plotted for selected significant differences in binding or activation. A Spider Web diagram was created using Microsoft Office Excel software. Figure design was adapted using Affinity Designer 2. Schematic images were created using BioRender software (BioRender.com; license holder: Katja Hoffmann).

## 3 Results

### 3.1 Establishment of the BW5147-FcγRζ reporter cell assay for CRP detection

The setup of the BW5147-FcγRζ reporter cell assay was adapted from our previously developed assays (20,33) and modified to analyze CRP-mediated activation of FcγRs (**Figure 1A**). BW5147-FcγRζ reporter cells (20) expressing human FcγRI (CD64), FcγRIIaH (CD32aH), FcγRIIaR (CD32aR), FcγRIIb (CD32b), FcγRIIIaV (CD16aV), FcγRIIIaF (CD16aF), FcγRIIIb (CD16b), and human CD99 as a negative control, were characterized for FcγR expression by flow cytometry using APC-coupled antibodies. All BW5147 cell lines expressed the transduced extracellular domain of the respective human FcγR or human CD99 (**Figure 1B**). The density of FcγRs expressed on the cell surface was largely comparable, but not identical, between different cell lines. As observed before, high-affinity BW5147-FcγRI (CD64) cells expressed lower amounts of FcγRs than transfectants expressing low-affinity FcγRs, i.e., CD32 and CD16, potentially due to the additional Ig-like domain (20).

**Figure 1:**
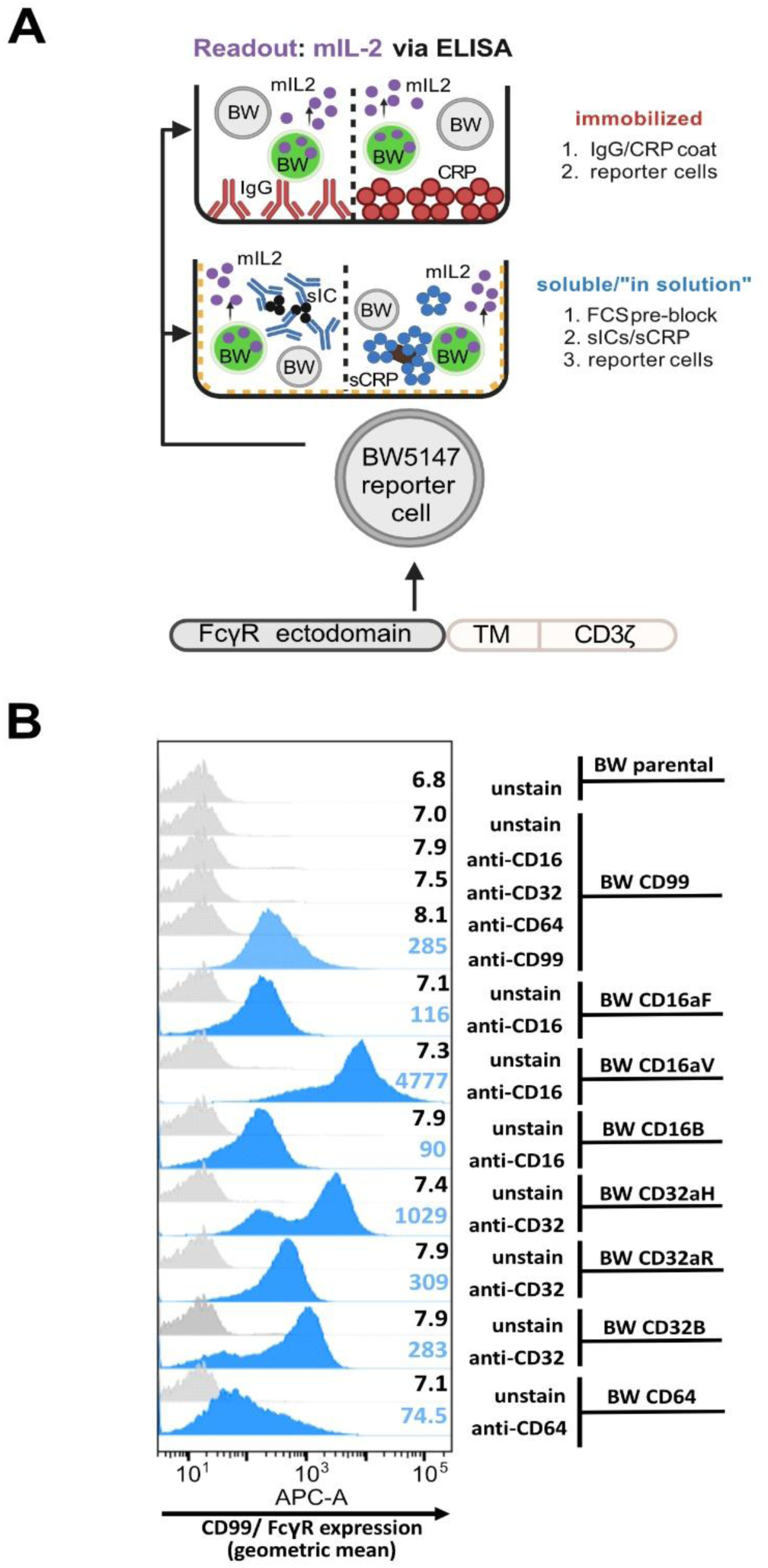
Setup of the BW5147-FcγRζ reporter cell assay and flow cytrometry-based analysis of FcγR expression: (**A**) Schematic of the assay setup: BW5147 cells stably express chimeric FcγR-CD3ζ-chain receptors leading to secretion of mIL-2 upon FcγR-activation, which can be mediated by immobilized human IgG or CRP as well as by soluble IgG-immune complexes (sol. ICs) or soluble CRP (alone or in complex with streptococci). The soluble assay setup requires pre-blocking of well plates with 10% FCS. mIL-2 levels in supernatant are measured by sandwich ELISA. (**B**) Characterization of BW reporter cells with anti-CD16/32/64-antibodies. A total of 100,000 BW5147 cells per sample were incubated with 100 µl flow cytrometry buffer containing a 1:100 dilution of the respective anti-CD-APC antibody for one hour on ice. Unstained BWCD99 cells and BWCD99 cells incubated with respective anti-CD antibodies served as negative controls. Additionally, BWCD99 cells were stained with anti-CD99-APC as positive control. Cells were analyzed by flow cytometry using a FACS Fortessa instrument and FlowJo software, gating on the main population of living cells.

### 3.2 CRP-dependent crosslinking selectively activates BW5147 reporter cells expressing CD64 (*FcγRI*), BWCD32aR (*FcγRIIaR*) and BWCD32b (*FcγRIIb*), and pCRP* is the major mediator of FcγR triggering

FcγR activation occurs upon receptor crosslinking by specific ligands. This is achieved either by immobilized or by soluble multimeric FcγR ligands, e.g., IgG immune complexes (20,33). Accordingly, immobilization of human IgG on MaxiSorp plates is the most basic BW5147-FcγRζ assay format (**Figure 1A**). This setup was transferred to pCRP by its immobilization on MaxiSorp wells at graded concentrations. As reported previously, all reporter cell lines became consistently activated when exposed to immobilized human IgG1 (**Figure 2A, upper panel**) (20,33). In contrast, only BWCD32aR (FcγRIIaR), BWCD32b (FcγRIIb) and BWCD64 (FcγRI) responded to immobilized pCRP, whereas we saw broad unresponsiveness in BWCD16aF (FcγRIIIaF), BWCD16aV (FcγRIIIaV), BWCD16b (FcγRIIIb) and BWCD32aH (FcγRIIaH) reporter cells (**Figure 2A, middle panel**). When experiments were jointly analyzed (AUC of activation after normalization to mean OD of individual experiments) pCRP-mediated activation was significant for BWCD32aR (FcγRIIaR; p<0.001) and BWCD64 (FcγRI; p<0.001) cells, whereas activation of BWCD32b (FcγIIb) was clearly detectable and reproducible but did not reach significance in two-way ANOVA/Dunnett’s multiple comparisons of all three ligands investigated (p=0.212) (**Figure 2B**). However, for individual analysis of pCRP as an activating ligand (one-way ANOVA/Dunnett’s multiple comparisons) BWCD32b (FcγIIb) activation was significant compared to the negative control (p=0.041). The limit of detection for pCRP was in the nanomolar range. Activation was dose-dependent for both IgG and pCRP, respectively, but responses induced by pCRP tended to be lower than those to IgG, except for high-affinity BWCD64 cells where AUCs were similar for IgG1- and CRP-mediated activation (**Figure 2B**). AUCs were significantly higher for all BW cell lines for IgG compared to negative control (‘no BWs’). AUCs for IgG1-mediated activation were significantly higher than for pCRP-mediated activation for all cell lines except BWCD32aR (FcγRIIaR) and BWCD64 (FcγRI) (two-way Anova/Tukey’s multiple comparisons, significance levels not indicated within the graph due to space constraints). Responses caused by pCRP-mediated activation were 60-70% of maximal IgG-mediated activation for BWCD32aR and BWCD32b cell lines and about 95% for BWCD64 cells (**Figure 2B**). Levels of CRP-induced mIL-2 responses did not correlate with surface expression levels of FcγR on BW5147 reporter cells, i.e. comparatively low levels of FcγRI were sufficient for higher activation levels than seen with CD32aR (FcγRIIaR), and CD32b (FcγRIIb). Strikingly, activation of BWCD32a (FcγRIIa) cells strictly depended on the allelic variant, with robust CRP-mediated activation of BWCD32aR (FcγRIIaR) cells, but no response in BWCD32aH (FcγRIIaH). This binary functional difference is remarkable as the variants differ only in one amino acid at position 131. Longer titrations for selected reporter cell lines and inclusion of BWCD99 cells as a negative control are shown in **Supplementary Figure 1A/B**.

**Figure 2:**
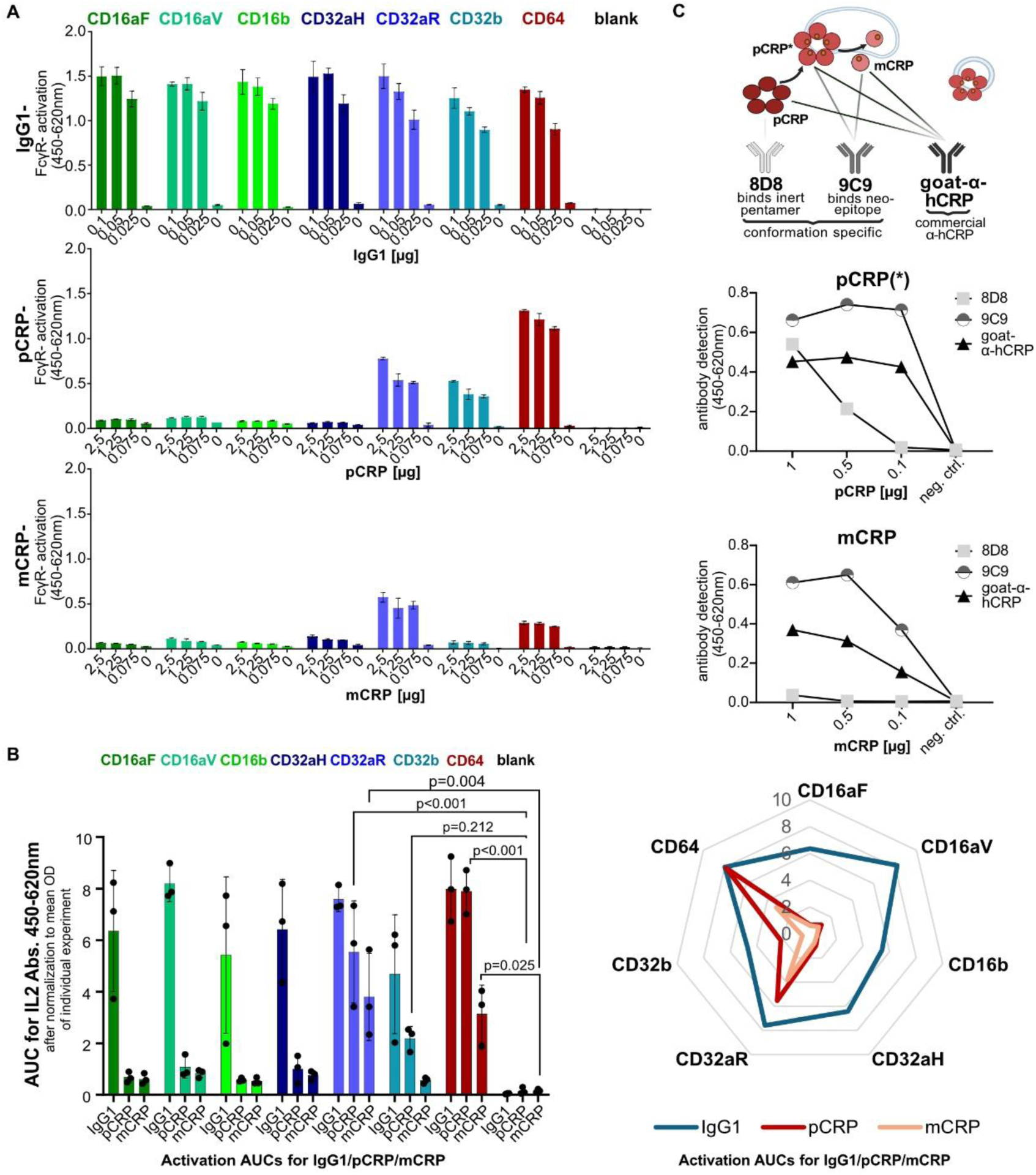
BW5147-FcγRζ reporter cell activation on immobilized IgG, pCRP and mCRP and binding of conformation specific antibodies to immobilized CRP isoforms: (**A**) <u>Upper:</u> mIL-2 levels produced by BW5147 cells on immobilized IgG1 (titrated from 0.1 to 0.025 µg in 50 µl medium). Each cell line was stably transduced with one FcγR only. Cell medium without BW cells (‘blank’) served as a negative control. <u>Center/lower:</u> mIL-2-levels produced by BW5147 reporter cells on immobilized pCRP (center) and mCRP (lower) (titrated from 2.5 to 0.63 µg in 50 µl medium, with concentrations matched using Qubit Fluorometric Quantitation). A total of 100,000 BW 5147 reporter cells were added to each well in 200 µl RPMI BW medium and incubated overnight at 37°C 5% CO_2_. Data are shown in technical replicates (N=3) with standard deviation for one representative of at least three individual experiments for each cell line. Activation shown as OD in a sandwich mIL-2-ELISA. (**B**) <u>Left:</u> AUCs for activation of BW cells by immobilized IgG1, pCRP and mCRP after normalization of ODs to mean OD of individual experiment. AUCs were calculated and jointly analyzed for three independent experiments normalized to mean OD of individual experiment with three technical replicates each. Two-way ANOVA and Dunnett’s multiple comparisons calculated using GraphPad Prism software. <u>Right:</u> Spider web plot of AUCs normalized to mean OD of individual experiment, created using Microsoft Excel Graph Software. (**C**) <u>Upper</u>: Schematic indicating recognition by conformation-specific monoclonal and polyclonal anti-CRP antibodies. Middle<u>/lower</u>: pCRP/mCRP was titrated from 1 µg to 0.1 µg/well and coated to MaxiSorp wells. Concentrations of pCRP and mCRP preparations were matched using Qubit Fluorometric Quantitation. CRP was detected using conformation specific 8D8 (anti-pCRP), 9C9 (anti-mCRP/pCRP*) and polyclonal goat anti-hCRP antibody.

There is evidence that the native CRP pentamer undergoes conformational changes before its ultimate dissociation into monomeric CRP (10). CRP isoforms were found to differ in their interaction with C1q, but very little is known about the functional impact of distinct CRP isoforms on single FcγR family members interaction (10,12,31,42,43). We then explored how our assay could be used to generate insights and new hypotheses on this issue (10–12,43). To find out more about conformational changes in the CRP pentamer induced by passive binding to MaxiSorp ELISA wells (designed for binding of medium to large sized hydrophilic biomolecules), an ELISA-based detection assay was performed using conformation-specific as well as polyclonal anti-CRP antibodies (**Figure 2C**). mCRP was generated as described previously (39) and concentrations of pCRP and mCRP preparations were matched using Qubit Fluorometric Quantitation. Conformation-specific monoclonal mouse anti-human CRP antibodies clone 8D8 (anti-pCRP) and clone 9C9 (anti-pCRP*/mCRP), and polyclonal goat anti-human CRP antibody (**Figure 2C, upper schematic**) were compared using graded concentrations of mCRP and pCRP preparations (12,43,44). All three CRP antibodies showed concentration-dependent binding to the coated pCRP (**Figure 2C, middle panel**), albeit with varying strength. mAb 8D8 recognized exclusively the inert pentamer exhibiting the weakest binding, particularly at low pCRP densities. As expected, 8D8 lost its capability to recognize CRP completely when tested with monomeric CRP (**Figure 2C, lower panel**). In contrast, mAb 9C9, which recognizes a neoepitope induced within the CRP pentamer and maintained after CRP fragmentation exhibited superior binding to mCRP, but also to pCRP. This finding indicates that the conformational change from pCRP to pCRP* has occurred to a relevant extent upon pCRP binding to the hydrophilic MaxiSorp surface. This observation is in line with the findings of Lv and Wang, who observed binding of both pCRP-specific and mCRP-specific antibodies upon immobilization on MaxiSorp plates and concluded that the dual antigenicity resulted from pCRP* expression rather than mixture of pCRP and mCRP (46).

As mCRP exposes the 9C9-defined neoepitope uncovered in pCRP* but has lost the 8D8-defined epitope characterizing native pCRP, we went on to investigate how activation is caused by coated mCRP compared to activation caused by coated pCRP in the BW5147-FcγRζ assay platform (**Figure 2A, lower panel**). Activation levels caused by mCRP were significantly diminished compared to pCRP for BWCD64 (FcγRI) cells (p=0.001, not depicted in the graph due to space constraints) and moderately diminished for BWCD32aR (FcγRIIaR) cells (p=0.357, not depicted). BWCD32b (FcγRIIb) cells showed minimal activation on coated mCRP. To exclude the possibility that mCRP preparations harmed the BW5147 reporter cells, the same amounts (20+20 µg; 10+10 µg; 5+5 µg) of mCRP and pCRP were coated together before testing BWCD64 reporter cells (**Supplementary Figure 1C**). mCRP did not appear to harm the cells but had little effect on upregulating activation caused by pCRP. We concluded that pCRP*, as defined by mAb 9C9, represents the major conformation of CRP that triggers FcγRs, while mCRP still causes activation at clearly lower levels.

Figure 2B gives an overview of the activation profiles induced by IgG1, pCRP and mCRP comparing activation levels (AUCs) for three independent experiments after normalization to the mean OD of each experiment, shown as a bar graph (right) and a spider web plot to illustrate the patterns generated (left). Generally, activation levels caused by IgG1 were higher than for pCRP > mCRP. Activation levels varied for all ligands depending on the FcγR composition of each BW5147 reporter cell type. Consistently high activation levels were seen for IgG1-mediated activation throughout all cell lines, whereas pCRP only activated BWCD64 (FcγRI), BWCD32aR (FcγRIIaR) and BWCD32b (FcγRIIb) cell lines with higher activation levels than for mCRP, which activated BWCD32aR>BWCD64>BWCD32b cells at relatively low levels.

### 3.3 FcγR activation is caused by CRP itself-but detecting CRP binding in ELISA does not correlate with FcγR activation

To ensure that the observed reporter cell activation was solely caused by CRP itself and not by another potentially activating factor present in the CRP preparation used (US Biological Life Sciences), which is generated from patient ascites/pleural fluid, we compared the previously used CRP composition with recombinant CRP produced in *E. coli* (US Biological Life Sciences) with respect to their activation efficacy of BWCD64 and CD32b reporter cells. Patterns of activation for recombinant CRP precisely mirrored those of CRP purified from human ascites, indicating that CRP is necessary and sufficient to cause activation of FcγRs (Figure 3A). To exclude any effect of other components, e.g., sodium azide, the CRP preparation purified from ascites/pleural fluid was dialyzed against PBS with Ca ^2+^/Mg^2+^ through a dialysis membrane overnight as described previously (39). Dialysis had no significant effect on activation levels, confirming that CRP itself was the cause of the activation (**Supplementary** Figure 1D). As previous studies have shown that biological effects attributed to CRP are actually caused by LPS contamination of recombinant CRP preparations (47), we further excluded any effect of LPS on the BW5147 reporter cell assay system (Figure 3B). To this end, we added graded EU endotoxin units of LPS to our ascites/pleural fluid purified CRP preparation and additionally tested the potential effect of LPS alone on our cells by adding graded EU units/ml to the culture medium of the BWCD64 cells in this assay. LPS had no effect on FcγR-activation responses.

**Figure 3:**
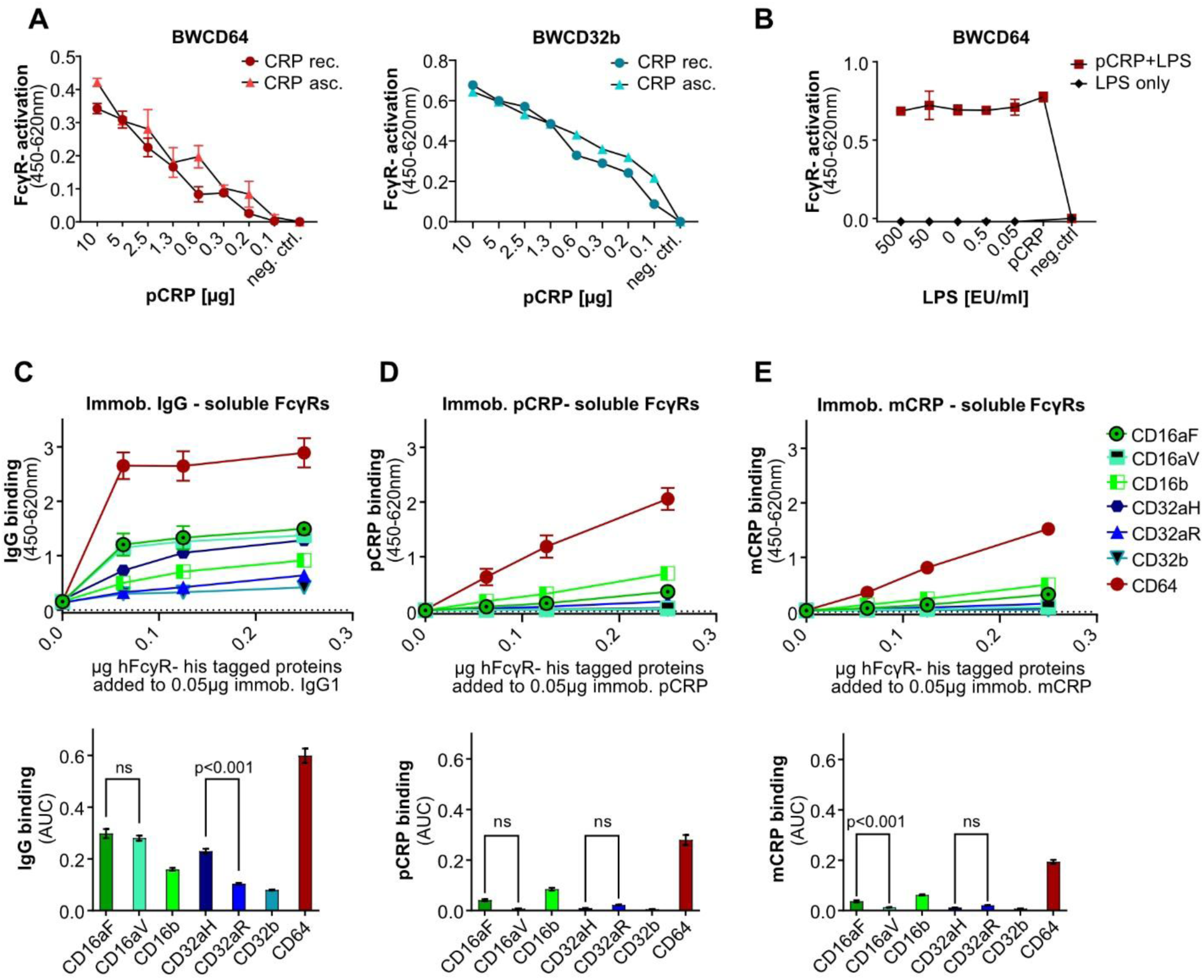
Effect of CRP source and addition of LPS on BW5147-FcγRζ activation and binding of epitope-tagged FcγRs to immobilized IgG, pCRP or mCRP: (**A**) BWCD64 and BWCD32b cell activation by recombinant human pCRP produced in *E. coli* and pCRP purified from human ascites/pleural fluid. pCRP was coated in graded amounts in PBS. A total of 100,000 BW5147 reporter cells were added to each well and incubated overnight. (**B**) Addition of graded amounts of LPS to a pCRP (5 µg/well) preparation was compared with activation caused by LPS only using BWCD64 reporter cells. LPS was added at the concentrations stated. EU units as stated by supplier: 1 mg/ml=1×10^6 EU/ml. 100,000 BW5147 cells were added to each well and incubated overnight. FcγR-activation shown as OD in sandwich mIL-2-ELISA after subtraction of background. (**C**/**D/E**) Titration of recombinant His-tagged hFcγRs from 0.25 µg to 0 µg in 50 µl PBS; binding to 0.05 µg coated IgG1/pCRP or mCRP/well. ODs for 450-620 nm. Data shown with standard deviation for two individual experiments with three technical replicates each. Calculation of AUCs of the binding curves using GraphPad Prism software. AUC for N=6 with standard error. Ordinary one-way ANOVA and Tukeýs multiple comparisons test carried out using GraphPad Prism software and selected significances are indicated on the graph.

The BW5147-FcγRζ assay demonstrated activation of FcγRs CD32aR, CD32b and CD64, but not of CD16aF, CD16aV CD16b and CD32aH by pCRP. However, interaction of pCRP with CD16 as well as CD32aH has been previously reported (23,24). Lu et al. observed binding to CD16, as did Temming *et al*., who additionally proposed a potential role in enhancement of IgG-mediated FcγR-activation through the interaction with pCRP. Nevertheless, CD64 and CD32a are proposed as the major CRP interactors, with a long-standing debate about the relevance of the CD32a allelic variants for both binding and activation (23,24,48–50). Since our assay system allows for unambiguously attributable responses of individual FcγRs and the available data on binding and activation were controversial, we set out to differentiate CRP-binding by and CRP-dependent activation of FcγRs as obtained in a comparable experimental setup.

To this end, the FcγR binding pattern to immobilized IgG1 (Figure 3C) was compared to immobilized pCRP (Figure 3D) and mCRP (Figure 3E) in a setup analogous to the activation setup, i.e. coated IgG1 and CRP and recombinant FcyRs added in solution for binding. AUCs for the individual binding curves were calculated (Figure 3C**/D/E, lower panels**). All recombinant his-tagged FcγR molecules showed binding to IgG1. Interestingly and in accordance with literature, the binding pattern observed for IgG1 (CD64>CD16aF/V>CD32aH>CD16b>CD32aR/CD32b) was different to the one of pCRP (CD64>CD16b>CD16aF>CD32aR>CD16aV/CD32aH/CD32b). The binding pattern for mCRP was similar to that of pCRP with slightly lower ODs. For IgG, in accordance with literature (51–53) binding for both allelic variants of CD32a was clearly detectable, with higher binding of the CD32aH allelic variant. The OD values measured in ELISA for IgG1-binding to the tested FcγRs were significantly higher than for pCRP. This observation correlates with the activation levels observed in the BW5147-FcγRζ activation assay and generally supports lower affinities of FcγR for pCRP. All human FcγRs bound to pCRP and mCRP with relatively lower strength as indicated by lower OD values. Binding of pCRP to CD64 showed the highest ODs/AUC, followed by CD16b>CD16aF>CD32aR>CD16aV/CD32aH/CD32b. At a generally low level, CRP binding to the CD32aR allelic variant was higher than to CD32aH, but this difference did not reach significance. Notably, ELISA binding in this very comparable setup did not correlate with FcγR triggering in the BW5147-FcγRζ reporter assay. E.g., pCRP-binding of CD16b was clearly stronger than binding of CD32b. However, BWCD16b (FcγRIIIb) cells were not activated by pCRP, whereas pCRP did readily activate BWCD32b (FcγRIIb) cells (Figure 2A). Thus, binding as detected by ELISA seems to be necessary but not sufficient for FcγR triggering.

### 3.4 Divergent FcγR activation profiles of solution-phase ICs and CRP, with pCRP* as the key activating isoform

The preparation (“blocking”) of hydrophilic MaxiSorp surfaces with saturating amounts of serum proteins allowed us to detect multimeric immune complexes in solution (sICs) as activating ligands of certain FcγRs (20,37). To investigate whether unbound pCRP in solution phase is also capable of cross-linking FcγRs, the protocol established for sICs was adapted for use in a pCRP context. (i) Synthetic sICs, (ii) soluble pCRP and (iii) pCRP-*Streptococcus pneumoniae* complexes (39), respectively, were added to serum-blocked ELISA wells (Figure 4A). For the third approach, to allow ligand binding to the B-face of the molecule and promoting the formation of pCRP*, pCRP was gpre-incubated with *Streptococcus pneumoniae serotype 27* containing high amounts of ‘C’-cell-wall-polysaccharide (CWPS). As observed before (20), soluble ICs formed by recombinant antigen and a recombinant monoclonal antibody, i.e., TNFα trimers and Infliximab, efficiently activated BWCD16aV (FcγRIIIaV) and BWCD32b (FcγRIIb) but neither BWCD32aR (FcγRIIaR) nor BWCD64 (FcγRI) reporter cells (Figure 4A**, upper panel**). In contrast, pCRP in solution phase and pCRP pre-bound to streptococci activated BWCD64>BWCD32aR>BWCD32b>BWCD16aV reporter cells, with BWCD16aV (FcγRIIIaV) and BWCD32b (FcγRIIb) only being slightly activated (Figure 4A**, middle and lower panel**). Therefore, the FcγRs that can be activated by pCRP and sICs in the solution phase are clearly distinct. Overall activation levels induced by sICs were higher than those caused by soluble CRP. Levels of BWCD64 (FcγRI) activation were substantially higher after pre-incubation with streptococci. This trend was less pronounced for BWCD32aR (FcγRIIaR) and BWCD32b (FcγRIIb) (Figure 4A**, lower panel**). Subsequently, activation levels induced by immobilized pCRP were compared with those induced by pCRP in solution or in solution pre-incubated with streptococci for recognition by BWCD64 (FcγRI) reporter cells (Figure 4B). As observed before, pre-incubation of pCRP with streptococci upregulated activation levels as compared to soluble pCRP only. This effect reached significance for 5 µg pCRP (p=0.044), but not 10 µg and 20 µg of pCRP (p=0.102 and p=0.204, respectively, Two-way ANOVA and Tukey’s multiple comparisons test). However, the activation induced by immobilized pCRP on MaxiSorp surfaces was significantly higher than both solution-phase approaches (with and without pre-incubation with streptococci) for all pCRP concentrations investigated (not all p-values are indicated on the graph for space constraints, p-values for 20 µg pCRP: p=0.005 and p=0.033 for comparison of immobilized pCRP-mediated activation to soluble pCRP only and soluble pCRP pre-incubated with streptococci, respectively). As pre-incubation with streptococci as well as immobilization on well surface both favor pCRP* conformation these results support pCRP* likely being the FcγR activating CRP conformation.

**Figure 4:**
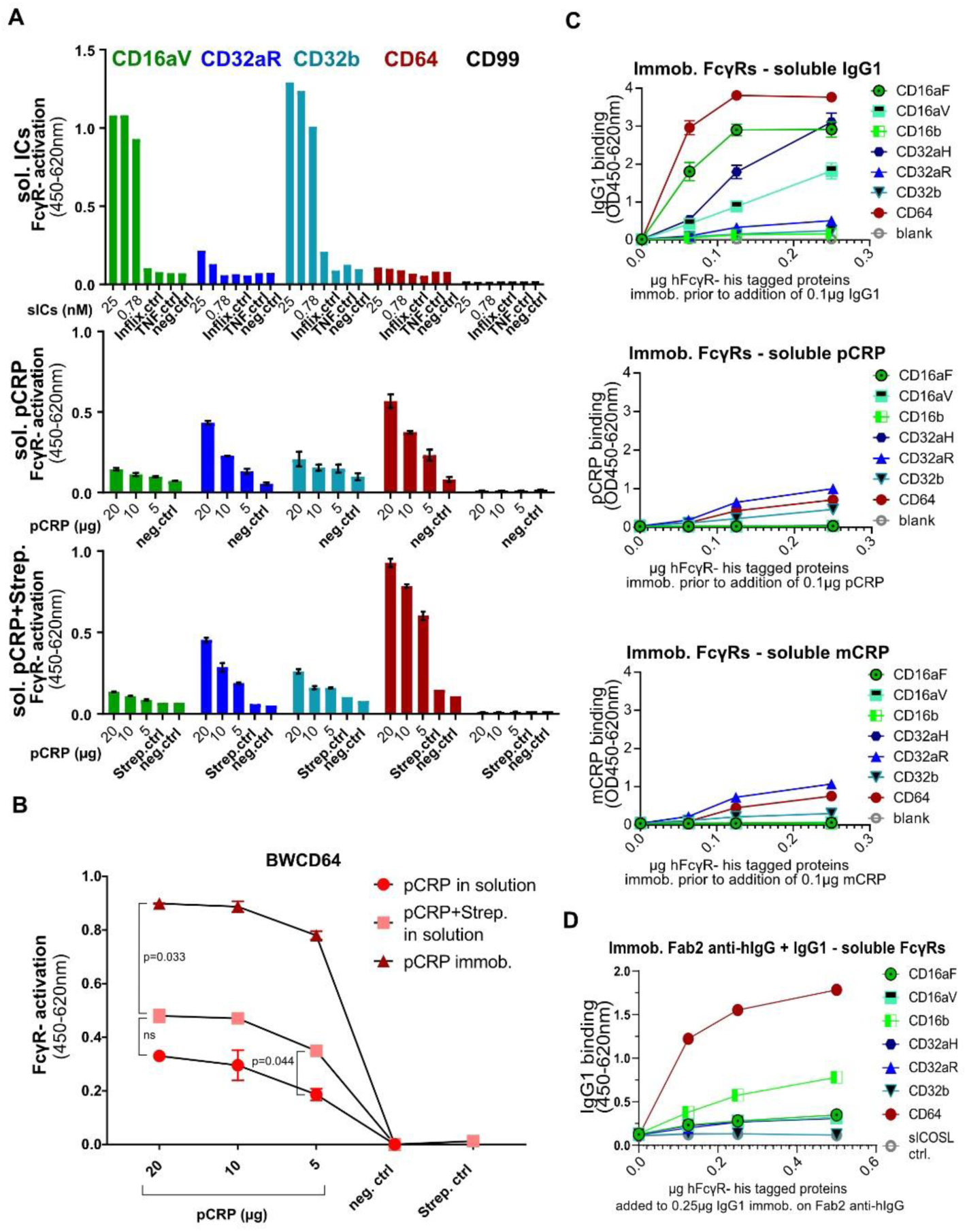
In solution phase BW5147-FcγRζ reporter assay for sol. ICs, sol. pCRP and sol. pCRP-streptococci complexes and binding of soluble IgG, pCRP and mCRP to coated His-tagged FcγRs: (**A**) MaxiSorp ELISA plates were saturated with 10% FCS. sICs as well as soluble CRP-streptococci complexes (with *S. pneumoniae* serotype 27) were allowed to incubate for two hours at RT prior to adding them to the experiment. <u>Upper</u>: sICs were added in 100 µl/well medium and consisted of 25 nM Infliximab (149.1 kDa) and 50 nM TNFα monomer (17.5 kDa) to ensure 1:1 stoichiometry (per ml stock of 25 nM ICs: 0.875 µg TNFα + 2.66 µg Infliximab). Selected values of log2 titration depicted in this graph. <u>Central:</u> pCRP in solution assay without pre-incubation with streptococci. CRP was added in 100 µl medium. <u>Lower:</u> 10 µl of streptococci were added to 20/10/5 µg of CRP. Complexes were added to wells in 100 µl medium. 100,000 BW5147 reporter cells were added to each well in another 100 µl of medium. Activation shown as OD in sandwich mIL-2-ELISA. Data are shown with standard deviation (N=2; N=1 for IC controls). (**B**) BWCD64 activation assay comparing coated pCRP and soluble pCRP/ soluble pCRP-streptococci complexes (N=2). Ordinary one-way ANOVA and Tukeýs multiple comparisons test carried out using GraphPad Prism software and selected significances are indicated on the graph. (**C**) Titration of His-tagged hFcγRs from 0.25 µg to 0 µg and coating to ELISA wells. Addition of 0.1 µg IgG1 (<u>upper</u>), pCRP (<u>central</u>) or mCRP (<u>lower</u>) and detection via goat-anti-hCRP antibody and DAG-POD for CRP and anti-human-IgG-POD for IgG1. ODs for 450-620 nm. Data shown with standard deviation for two individual experiments with three technical replicates each. (**D**) Coating of goat F(ab)_2_ anti-human IgG (Fab-specific) (0.1 µg in 50 µl/well) was followed by blocking and addition of hIgG1 (0.25 µg in 50 µl/well) before addition of soluble human FcyR-His-proteins titrated as stated in the graph. Detection with rabbit anti-His antibody and GAR-POD was performed. Data shown in technical triplicates for two individual experiments.

To investigate, whether the ‘in solution’ activation could be correlated with an ‘in solution’ binding approach of CRP to FcγRs, we reversed the setup of our binding assay established before (Figure 3C**/D/E**). After coating MaxiSorb wells with recombinant His-tagged human FcγR proteins, IgG1 (Figure 4C**, upper**), pCRP (Figure 4C**, center**) or mCRP (Figure 4C**, lower**) were added and binding was investigated through goat anti-hCRP/DAG-POD or anti-human-IgG-POD for CRP and IgG, respectively. AUCs are compared in **Supplementary** Figure 2. For IgG1, the pattern observed in this ‘reversed setup’ was widely comparable to the pattern in the initial binding assay (CD64>CD16aF>CD32aH>CD16aV>CD32aR>CD16b>CD32b). Binding of the CD32aH allelic variant was significantly higher than for the CD32aR allelic variant (p<0.001; One-way ANOVA and Tukeýs multiple comparisons for AUCs; **Supplementary** Figure 2), as observed previously, whereas – in contrast to published literature (51) - a stronger binding to the CD16aF than to the CD16aV allelic variant (p<0.001) was seen in this setup, suggesting that the experimental conditions of the chosen assay setup could influence the extent of binding. To compare the effect of presentation of binding partners, human IgG1 was immobilized using goat F(ab)_2_ anti-human IgG (Fab-specific) followed by the addition of soluble human FcγR-His-proteins (Figure 4D). Now CD16aV binding matched CD16aF binding. This confirms the impact of the presentation of the binding partners in these test formats and the need to compare different setups. Notably, for both pCRP and mCRP, the binding pattern obtained in this ‘reversed setup’ largely mirrored the activation pattern, with CD32aR, CD64 and CD32b showing the highest binding affinities (Figure 4C). Thus, although FcγR activation is generally strongest upon ligand immobilization, binding of soluble ligands to immobilized FcγR confirms the receptor crosslinking potential of ligands.

### 3.5 CRP ligand-ligand interactions in FcγR activation

Since pCRP, monomeric IgG, and soluble ICs represent distinct ligands that share the same immunological compartments, they may be recognized simultaneously by FcγRs. We applied the BW5147 reporter cell assay system to explore interactions between these ligands. First, the effect of pCRP in solution on activation caused by soluble ICs was investigated. As sICs were shown to efficiently activate BWCD16aV (FcγRIIIaV) and BWCD32b (FcγRIIb) cells (20), these reporter cell lines were chosen to investigate a possible inhibitory effect of pCRP. Two different concentrations of sICs (3 nM and 0.5 nM) were chosen to ensure that the effect of the addition of pCRP was analyzed under conditions of both high and low sIC-mediated activation. sICs were generated prior to addition of different concentrations of pCRP before adding BWCD16aV (FcγRIIIaV) or BWCD32b (FcγRIIb) reporter cells. pCRP in solution did not show any impact on sIC-mediated activation of both FcγRs tested, even when high concentrations of pCRP were added (Figure 5A).

**Figure 5:**
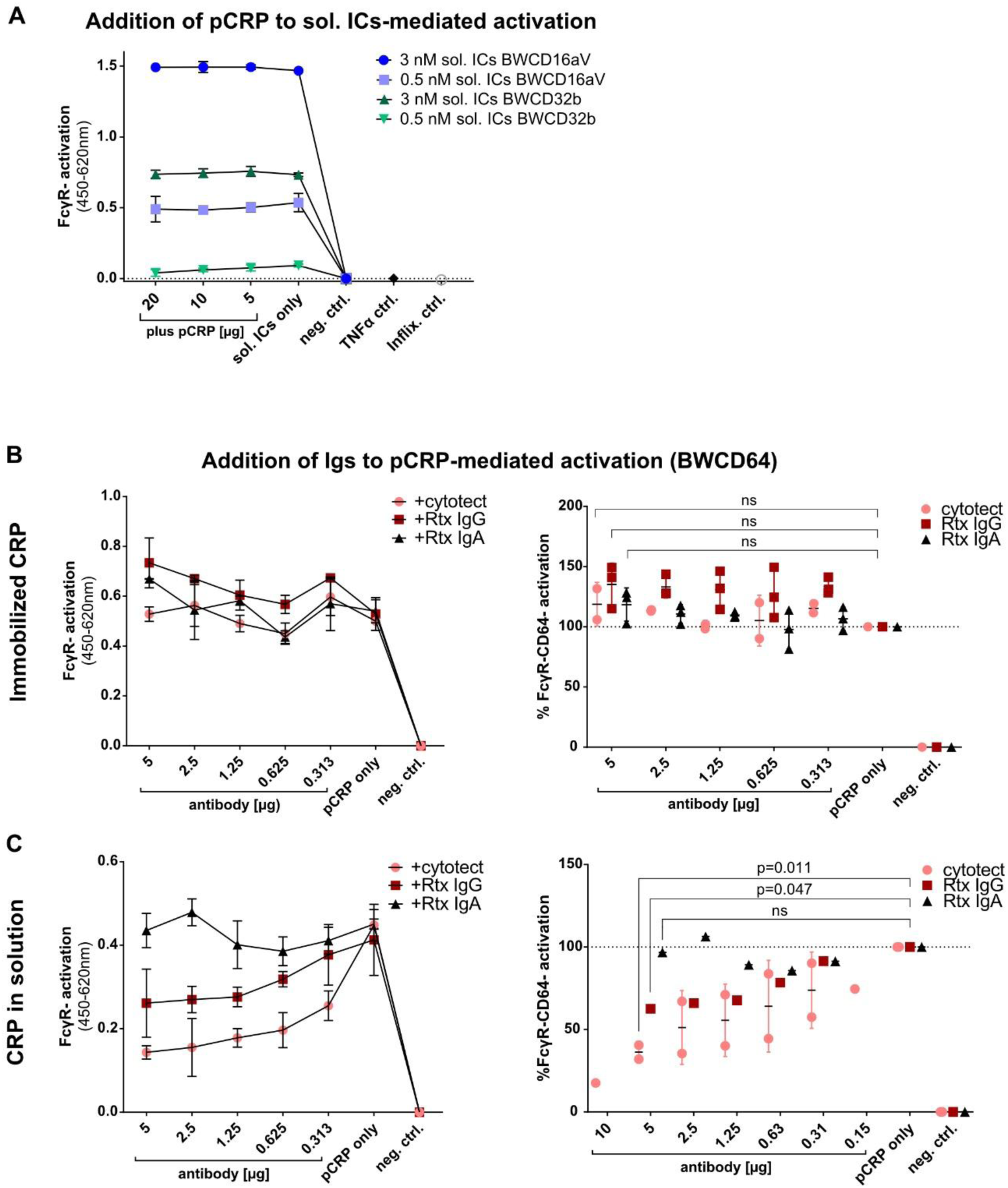
Competitive binding assays for distinct FcγR-ligands: (**A**) BW5147-FcγRζ reporter assay “in solution” with sol. ICs and added pCRP: ELISA wells were blocked with 10% FCS. Sol. ICs were allowed to incubate for one hour at RT in 100 µl BW medium prior to adding pCRP for 30 minutes. Both were added to 100 µl BW medium with 100,000 BWCD16aV or BWCD32b cells. sICs consisted of 3/0.5 nM Infliximab (149.1 kDa) and 6/1 nM TNFα monomer (17.5 kDa) to ensure 1:1 stoichiometry. Data are shown with standard deviation (N=2). Activation shown as OD in sandwich mIL-2-ELISA minus background control. **(B/C)** pCRP was immobilized (**B**) or added ‘in solution’ in 50 µl BW medium to pre-blocked wells (**C**). 5 to 0 µg immunoglobulins cytotect®, Rtx IgG or Rtx IgA were added in 100 µl (**B**)/ 50 µl (**C**) BW medium 15 minutes prior to addition of 100,000 BWCD64 cells in 100 µl medium. Representative individual experiments in technical replicates (N=2) are shown on the <u>left</u>. The <u>right</u> side summarizes three/two independent experiments for activation caused by immobilized pCRP (setup **B**) or soluble pCRP (setup **C**) after normalization to “CRP only”. Activation is caused by 10 µg coated or 15 µg soluble pCRP per well, respectively. Ordinary one-way ANOVA and Tukeýs multiple comparisons test carried out using GraphPad Prism software for 5 µg antibody results compared to “CRP only” control.

**Figure 6:**
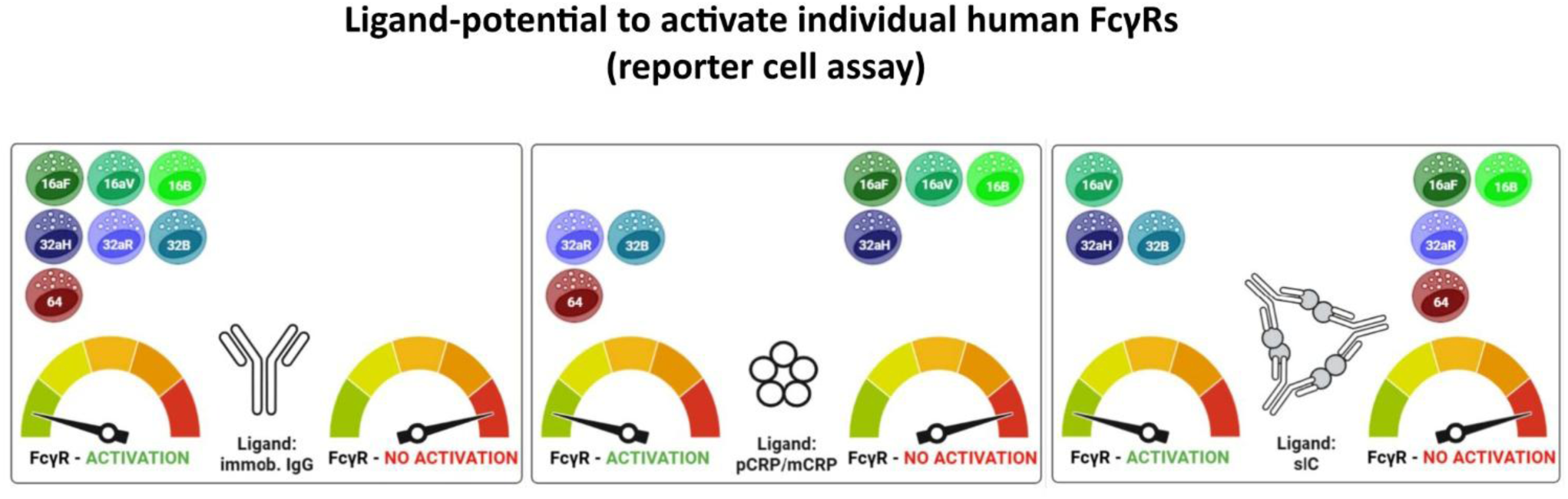
Graphical abstract summarizing FcγR-activation patterns for IgG, p/mCRP and soluble ICs.

Next, we investigated the impact of soluble, monomeric IgG on activation caused by (i) immobilized pCRP or (ii) unbound pCRP in solution phase. For this pCRP was coated (Figure 5B) or added to FCS-pretreated MaxiSorp plates keeping pCRP in solution (Figure 5C) before graded amounts of purified polyclonal human IgG (cytotect®), monoclonal Rituximab IgG1 (Rtx) or monoclonal Rtx IgA as control were added. None of these immunoglobulins caused a decrease in BWCD64 (FcγRI) activation levels mediated by immobilized pCRP (Figure 5B). A slight, non-significant increase in activation, especially seen with Rtx IgG1 (p=0.066), is likely attributable to residual Rtx binding after blocking of CRP-coated ELISA wells. However, when testing the activation caused by pCRP in solution, cytotect® caused a significant (p=0.011 for 5µg), dose-dependent decrease in BWCD64 (FcγRI) activation (Figure 5C). The addition of Rtx IgG1 also caused a significant decrease in activation levels (p=0.047), though this effect was less pronounced than for polyclonal IgG in cytotect®. Activation levels caused by solution-phase pCRP supplemented with Rtx IgA as a control remained unaffected (Figure 5C).

## 4 Discussion

The role of CRP as an activating ligand of FcγRs has been a matter of debate for decades (21–27,48). Unraveling the bimolecular interactions between FcγRs and CRP is complicated by a number of intricate details. The confusion stems from complex experimental settings with different readouts, but also from the variety of existing FcγRs, different types of immune cells expressing different ranges of FcγRs, the influence of FcγR ligands other than CRP, the variability of CRP preparations, and finally CRP itself, which acquires intermediate conformations and isoforms, including pCRP, pCRP*, and mCRP, with different biophysical properties and functional consequences. Experimental setups using antibody-based detection systems are delicate because they may affect the Fc-binding capacity of human FcγRs, and there is potential for antibody-species cross-reactivity (e.g., binding of mouse IgG (54)). Furthermore, the binding of CRP to human FcγRs is low-affinity and, as an opsonin with less binding specificity than immunoglobulins (5,55), CRP interacts with many pathogen- or damage-associated patterns (e.g. apoptotic cells, oxLDL, phosphocholine groups (55,56)) that may be present in the reagents used, making low affine binding of FcγRs to CRP even more difficult to detect. Moreover, CRP and CRP-bound molecules interact with multiple cellular receptors, such as different FcγRs. In addition, CRP-mediated amplification of TLR signaling (27) complicates the attribution of the resulting signaling cascades in cells. In this situation, a highly reductionist assay approach, as explored here, is essential to analyze and quantify CRP-mediated molecularly defined interactions with individual FcγRs leading to receptor cross-linking.

### 4.1 Future applications of the BW5147-FcγRζ reporter cell assay

The BW5147-FcγRζ reporter cell panel allows rapid screening of FcγR types and isoforms and their discrimination into CRP-receptive, CRP-unresponsive and decoy FcγRs. Key advances of this reporter system include high accuracy and hierarchical resolution of FcγR type-specific activation compared to traditional indirect assessments such as CRP binding, and a scalable and quantifiable methodology that provides flexible high-throughput readouts such as mouse IL-2 detection in cell culture supernatants or CD69 plasma membrane densities (20). FcγR profiling and classification have important implications for a better understanding of pCRP in immune defense, inflammation, and autoimmune disorders. The incorporation of Fc-less Fab fragments from conformation-dependent CRP-specific monoclonal antibodies (12,44,45) in the BW5147-FcγRζ assay may allow more precise conclusions to be drawn about the intramolecular steps that ultimately lead to FcγR cross-linking. This approach may provide further insight into the molecular sequence of events leading from native pCRP to pCRP* to its degradation and finally to mCRP, which was found to be less efficient in activating FcγRs. Likewise, pharmaceutical CRP inhibitors such as phosphocholine mimetics (40), and physiological modulators like Ca^2+^ and C1q (23,57) can be analyzed and screening for new drugs with superior efficacy will be possible. Although it has unique advantages, the reduction of an experimental set-up to a certain “necessary minimum” raises the question of the relevance of factors not taken into account. The relevance of this fact can be seen in cases where CRP does not exert a function alone, but rather affects the interplay of different ligands and receptors, e.g., enhancing the activation caused via TLRs (27) or in the interaction of FcγRs and C5a-receptor (58). This might be relevant for CRP-mediated activation of non-classical, CD16-positive monocytes and interaction of CRP isoforms with NK cells. The lack of CRP-mediated activation of CD16 isoforms in our reductionistic setup raises the question of potential co-receptors, like CD88/C5aR1 needed for activation or dependency on lipid rafts (59,60). While not all scenarios can be addressed in our setup, in cases where CRP co-engages with receptors, co-expression of such immune receptors by BW5147-FcγRζ reporter cells may be feasible in the future.

### 4.2 CRP profiles of individual FcγRs as revealed by the BW5147-FcγRζ reporter cell assay

The BW5147-FcγRζ reporter assay allows for individual exploration of CRP-FcγR interaction resulting in effective receptor crosslinking rather than simple CRP binding. In agreement with the literature (21–24,27), we observed a readily induced activation of CD64 (*FcγRI)* and FcγRs CD32a and CD32b (*FcγRII*) but not CD16aF, CD16aV, and CD16b (*FcγRIII)*. Pronounced differences were noted concerning FcγRII/CD32. The inhibitory FcγR CD32b as well as the activating allelic variant CD32aR responded to pCRP, but CD32aH did not. Again, this finding confirms earlier reports (23,24,26,27). This allelic restriction was also observed in the ‘reverse’ ELISA binding assay, where CD32aR binding to coated CRP was in clear contrast to the very slight binding to CD32aH (Figure 4, less pronounced in the ‘non-reversed’ setup of Figure 3). Even though CRP binding to CD16aF or CD16b appeared stronger than CD32b in the ‘standard’ ELISA binding assay, no receptor crosslinking could be detected in this setting, whereas the FcγRs showing the highest binding affinity to CRP in the ‘reverse’ setup - CD32aR, CD64 and CD32b - were readily activated in the BW5147-FcγRζ reporter cell assay. Thus, the ‘in solution’ binding potential might be indicative of subsequent FcγR activation in this setting.

### 4.3 Insights into CRP conformation causing FcγR activation

While binding to and activation of FcγRs by CRP has been reported, little is known about the conformational isoforms of CRP that are capable of triggering FcγRs. Binding studies have analyzed the interaction of non-ligand-bound pCRP (23,24), whereas experimental systems of CRP-mediated FcγR activation have often used ligand-bound CRP after pre-incubation with CWPS (27), streptococci (26) or Zymosan (23). Although binding to these ligands favors the formation of pCRP* conformation, as has been described for binding to PC groups on activated cell membranes and microvesicles (10), the conformation of the CRP isoform(s) that cause activation of individual FcγRs remains elusive.

Here, we present several lines of observation pointing to pCRP* as the major FcγR-activating CRP isoform. First, compared to pCRP in the solution phase, FcγR activation was significantly higher for immobilized pCRP on hydrophilic MaxiSorp surfaces. Second, pre-incubation with streptococci, likely favoring pCRP* conformation, increased activation levels compared to soluble pCRP. Third, activation levels caused by immobilized pCRP were higher than for immobilized mCRP. Intriguingly, conformation-specific mAbs, i.e., anti-pCRP antibody clone 8D8 binding the inert pentamer and anti-pCRP*/mCRP (‘*neoepitope*’) antibody clone 9C9, revealed the simultaneous presence of both isoforms after coating of pCRP to MaxiSorp wells. The coating of MaxiSorp surfaces with mCRP confirmed exclusive recognition by mAb 9C9 at comparatively low levels of FcγR triggering (Figure 2C), supporting the notion that 9C9 reactive pCRP* is closely associated with FcγR activation. This notion is consistent with the findings of Lv and Wang, who compared binding pattern of pCRP-as well as mCRP/pCRP*-specific mAbs upon immobilization on hydrophilic MaxiSorp plates (46). Considering the plastic surface properties, the time course of the coating inducing conformational changes and the lack of binding of soluble pCRP by immobilized mCRP, they concluded that the dual mAb antigenicity of 9C9 and 8D8 is caused by pCRP* rather than mixture of pCRP and mCRP (46). Thus, surface immobilization on plastic surfaces is suggested as a simple way to generate pCRP* *in vitro*, mimicking the process that takes place on cell membranes *in vivo* (46). Contrary to mCRP, Lv and Wang found surface immobilized pCRP* to bind solution phase pCRP. As the amounts of CRP used in our studies were about ten times higher than those employed in binding studies by Lv and Wang, association of native pCRP molecules in solution phase to coated pCRP* could have occurred during our coating process, explaining why 8D8 reactive material is found upon immobilization of higher amounts of CRP to plates. This resembles *in vivo* scenarios on cell membranes or pathogen interfaces, with native pCRP molecules changing conformation towards the pCRP* isotype upon binding and subsequently facilitating the recruitment of further pCRP molecules. This concept reflects the coordination of both, the opsonic activity of pCRP* followed by effective crosslinking of FcγRs as essential mediators of phagocytosis.

Increasing evidence indicates that mCRP is initiating most pro-inflammatory actions of CRP as highlighted by its increased binding capacity to C1q and exposure of the cholesterol-binding sequence (7,61). Accordingly, mCRP is considered the relevant isoform of CRP in the regulation of local inflammation. Our findings could extend this concept: the data suggest that pCRP/pCRP* is generally also capable of mediating relevant immune effector functions via FcγR-bearing cells, and that mCRP-mediated activation of FcγRs is even lower than for pCRP*. In conjunction with the known differences in half-life between pCRP and mCRP the results suggest that CRP isoforms might trigger separate effectors, leading to step-by-step cascades of activation and decline.

### 4.4 Ligand-ligand interactions of CRP

IgG, CRP and soluble ICs are independently generated FcγR-ligands present within the same immunological compartments. This could allow for competitive binding and ligand displacement. We investigated the impact of these three ligands on activation mediated by any other one of the three. Interestingly, pCRP in the solution phase could not reduce the dominant activating effect of sICs on FcγRIIb/IIIaV (Figure 5A). However, conversely, pCRP-dependent activation of FcγRI showed significant inhibition by monomeric IgG but not by IgA used as a control. Intriguingly, the IgG levels employed in our experiments were in the range of 2.5 mg/dl which is at least one order of magnitude lower than the normal range of IgG levels in human serum (407-2,170 mg/dl) (62). Consequently, IgG may inhibit native pCRP-mediated activation even more pronouncedly than was documented in our experimental setting. Immobilization on MaxiSorp ELISA plates, however, enabled pCRP to activate FcγRI even in the presence of high concentrations of monomeric IgG (Figure 5B). Again, this observation along with the fact that *Streptococcus pneumoniae* serotype 27 increased pCRP bioactivity in the solution phase supports the concept that the conformational change of pCRP into pCRP* strongly increases its potency to crosslink FcγRs.

Notably, in the presence of physiological IgG concentrations found in plasma soluble pCRP has only negligible FcγRI/CD64 activating capabilities, implying an important anti-inflammatory role of IgG on CRP-dependent FcγR activation. In contrast, locally immobilized pCRP in a pCRP* conformation readily acquires FcγRI/CD64 activating capabilities, unaffected by the presence of monomeric IgG. The findings highlight the role of pCRP* for FcγR activation in localized inflammatory processes.

### 4.5 Limitations of the BW5147 FcγRζ reporter cell-based CRP detection

Limitations of the assay platform are evident when testing native human material, e.g., sera and other liquid specimens containing a variety of different proteins and immunoglobulins present. Additionally, for solution-phase examinations we only observed FcγR activation by pCRP when using relatively high concentrations of pCRP (>100 µg/ml), which are only present in patients under certain conditions, e.g., severe inflammation or sepsis. Competing, abundant FcγR ligands with higher affinities than pCRP, like IgG or sICs, are consistently present in patient’s liquid biopsies. Favored by the fact that BW5147 cells are largely inert to human cytokines, the BW5147-FcγRζ reporter cell platform has been successfully applied and validated for the highly sensitive detection of virus-specific IgG in serum, sICs in patient and animal samples, and viral FcγR ligands (21,34,35,37,63,64). The presence of such ligands with high FcγR affinity in native clinical materials could easily cause confounding effects on BW5147-FcγRζ reporter cells, making the attribution of measured bioactivities to pCRP difficult or even impossible. Nevertheless, it will be useful in the future to carefully explore possible applications of the new test system by combining it with quantitative, highly sensitive CRP assays in patient-derived material or for clinical research purposes.

### 4.6 Conclusions

Key advances of this reporter cell system include (i) its high accuracy and resolution of FcγR type-specific activation, (ii) a scalable and quantifiable assay with flexible high-throughput readouts in the nanomolar range, (iii) a reporter system sensitive to CRP isoforms, (iv) a comprehensive panel including all human FcγRs, and (v) a test system that allows easy integration of additional FcγR ligands and modifiers of CRP-mediated activation. In practice, the platform is suitable for implementation in small or large screening setups in research laboratories. This reporter cell approach allows for future adaptations, as the FcγR-bearing reporter cells can be engineered with additional CRP interactors and alternative reporter modules to optimize the methodology for specific applications.

**Table 1:** FcγR activation profiles induced by distinct ligands: FcγR-activation observed in BW5147-FcγRζ reporter assay for coating (‘crosslinking’) of IgG or p/mCRP, as well as soluble ICs (TNFα+Infliximab) and soluble pCRP (+/- pre-incubation with *S. pneumoniae* serotype 27). Activation “**+”** is defined as OD (λ=450-620 nm) 0.3-2.0 higher than background level OD, threshold activation “**(+)”** is defined as OD=0.15-0.3 higher than background level, no activation “-“ as OD<0.15 higher than background. NA= no data available.

|  | CD16aF<br>(*FcγRIIIaF*) | CD16aV<br>(*FcγRIIIaV*) | CD16b<br>(*FcγRIIIb*) | CD32aH<br>(*FcγRIIaH*) | CD32aR<br>(*FcγRIIaR*) | CD32b<br>(*FcγRIIb*) | CD64<br>(*FcγRI*) |
| --- | --- | --- | --- | --- | --- | --- | --- |
| Immob.<br>IgG | + | + | + | + | + | + | + |
| Immob.<br>p/mCRP | - | - | - | - | + | + | + |
| Sol. ICs | NA | + | NA | + | (+) | + | - |
| Sol. pCRP | NA | - | NA | NA | + | (+) | + |

## 5 Conflict of Interest

The authors declare that the research was conducted in the absence of any commercial or financial relationships that could be construed as a potential conflict of interest.

## 6 Author Contributions

Conceptualization: KH, HH; Methodology: AH, JZ, SUE, JT, PK, KP, HH, KH; Investigation: AH, JS, KH; Funding acquisition: KH, HH; Supervision: KH, SUE, HH; Writing – original draft: AH, KH, HH; Writing – review & editing: AH, JZ, KH, HH.

## 7 Funding

This work was supported by the German Research foundation (DFG) through FOR2830, HE 2526/9-2 and “NaFoUniMedCovid19“ (FKZ: 01KX2021 - COVIM to H.H. Further support was received in personal grants to SUE from the German Research Foundation (DFG) DFG EI 866/9-1 and EI 866/10-1. AH was supported by a stipend of the Cusanuswerk. JS was supported by a stipend of the University of Freiburg according to the Landesgraduiertenförderungsgesetz.

## Acknowledgments

We are grateful to Dr. Mark van der Linden, National Reference Center for streptococci, Department of Medical Microbiology, University Hospital (RWTH), Aachen, for providing *S. pneumoniae*.

We thank Prof Lawrence A. Potempa, College of Pharmacy, Roosevelt University, Schaumburg, IL, USA for kindly providing monoclonal antibodies detecting specific CRP conformations (clone 8D8 and 9C9).

We are grateful to Sheena Kreuzaler for technical assistance and support for mCRP generation. We thank Verena Horner for her support with mCRP purification. Mona Wolf for technical assistance with cell culture.

We are grateful to Anne Halenius and Zsolt Ruzsics for profound project discussions.

We acknowledge support by the Open Access Publication Fund of the University of Freiburg.

## 12 Data Availability Statement

The raw data supporting the conclusions of this article will be made available by the authors, without undue reservation.

## Use of BioRender Images in Figure 1, 2 and 6

Created in BioRender. Hoffmann, K. (2025): https://BioRender.com/g97k4ey; https://BioRender.com/n08p187; https://BioRender.com/n6jkoc7; https://BioRender.com/lgmkfx9; https://BioRender.com/c5sm64s

## Notes

### Competing Interest Statement

The authors have declared no competing interest.

