## Supplemental Figures for "Human Fcγ-receptors selectively respond to C-reactive protein (CRP) isoforms"

### *Supplementary Material*

#### Supplementary Figure 1

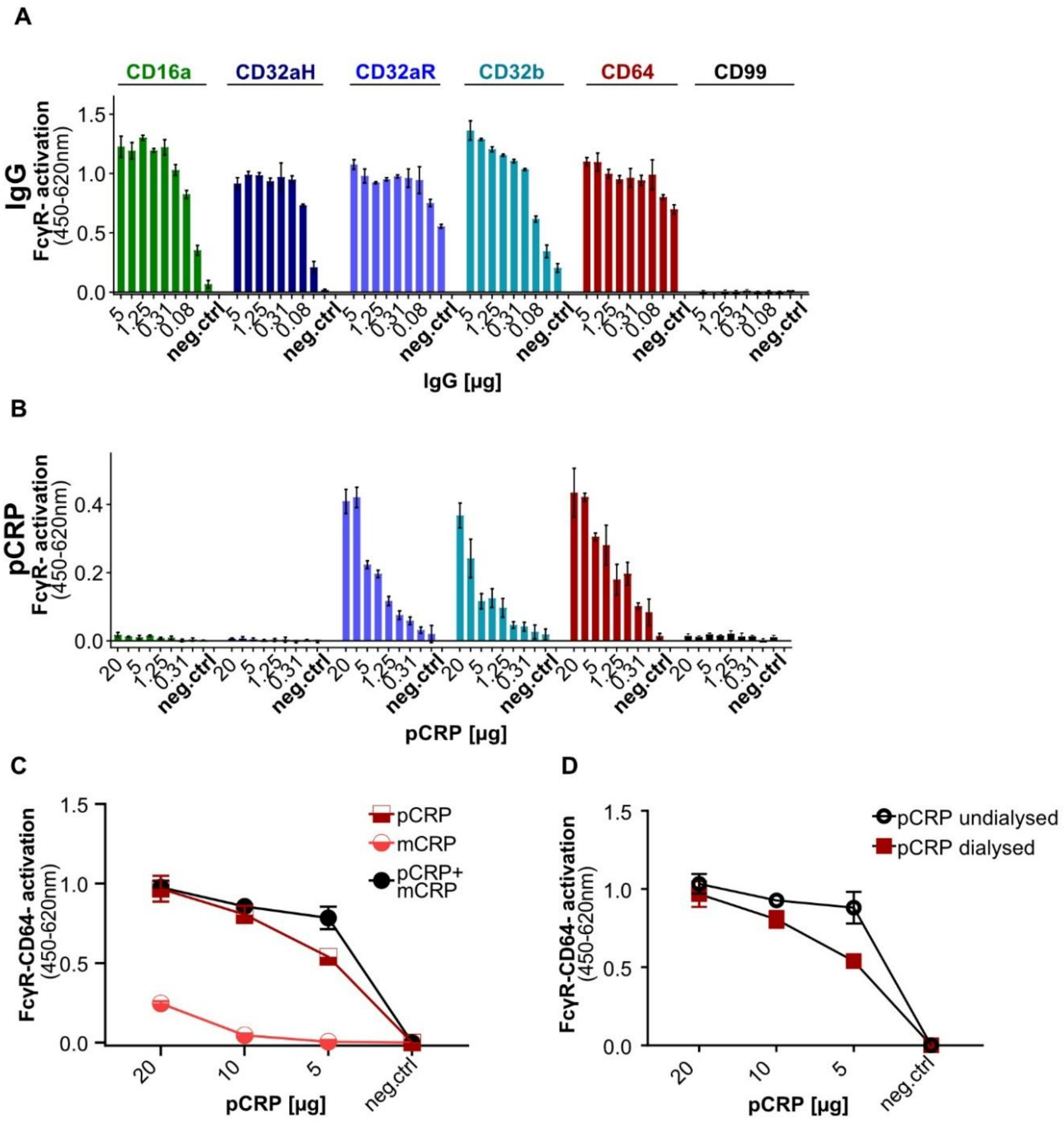


**Supplementary Figure 1:** Supplemental Figure 1: BW5147-FcγRζ reporter assay with longer titration and BWCD99 control, joint mCRP/pCRP activation for BWCD64 and effect of pCRP dialysis: (**A**) Activation of BW5147 cells on immobilized cytotect® as an IgG-source (titrated from 50 to 0,05 µg/ml in 100µl PBS). Each cell line was stably transduced with one FcγR only. BWCD99 cells served as a negative control. (**B**) Activation of BW5147 cells on immobilized pCRP (titrated from 200 to 0,2 µg/ml in 100µl PBS). 100.000 BW cells were added to each well in 200µl RPMI BW medium and incubated overnight at 37°C 5% CO2. Data shown in technical replicates (N=3) with standard deviation for one representative of at least three individual experiments for each cell line. Activation shown as OD in sandwich mIL2-ELISA. (**C**) BWCD64-activation caused by pCRP, mCRP or both coated together (each at the concentration stated). Concentrations of pCRP and mCRP preparations were matched using Qubit Fluorometric Quantitation. 100.000 BWCD64 cells were added to each well in 200 µl medium and incubated overnight at 37°C 5% CO2. Data shown in technical replicates (N=2) with standard deviation. (**D**) Comparison of activation in BWCD64 cell activation assay with equal amounts dialysed or undialysed CRP. 100.000 BW5147 cells were added to each well in 200 µl medium and incubated overnight at 37°C 5% CO2. Activation shown as OD in sandwich mIL2-ELISA after subtraction of background.

#### Supplementary Figure 2

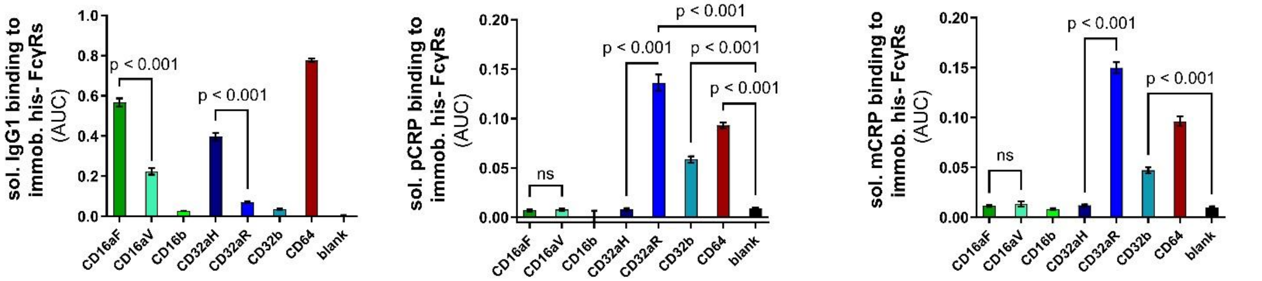


**Supplemental Figure 2:** AUCs for ‘in solution’ binding experiments: Calculation of AUC of the binding curves shown in Figure 4c using GraphPad Prism software. AUC for N=6 with standard error. Ordinary one-way ANOVA and Tukey´s multiple comparisons test carried out using GraphPad Prism software and selected significances are indicated on the graph.

#### Supplementary Figure 3

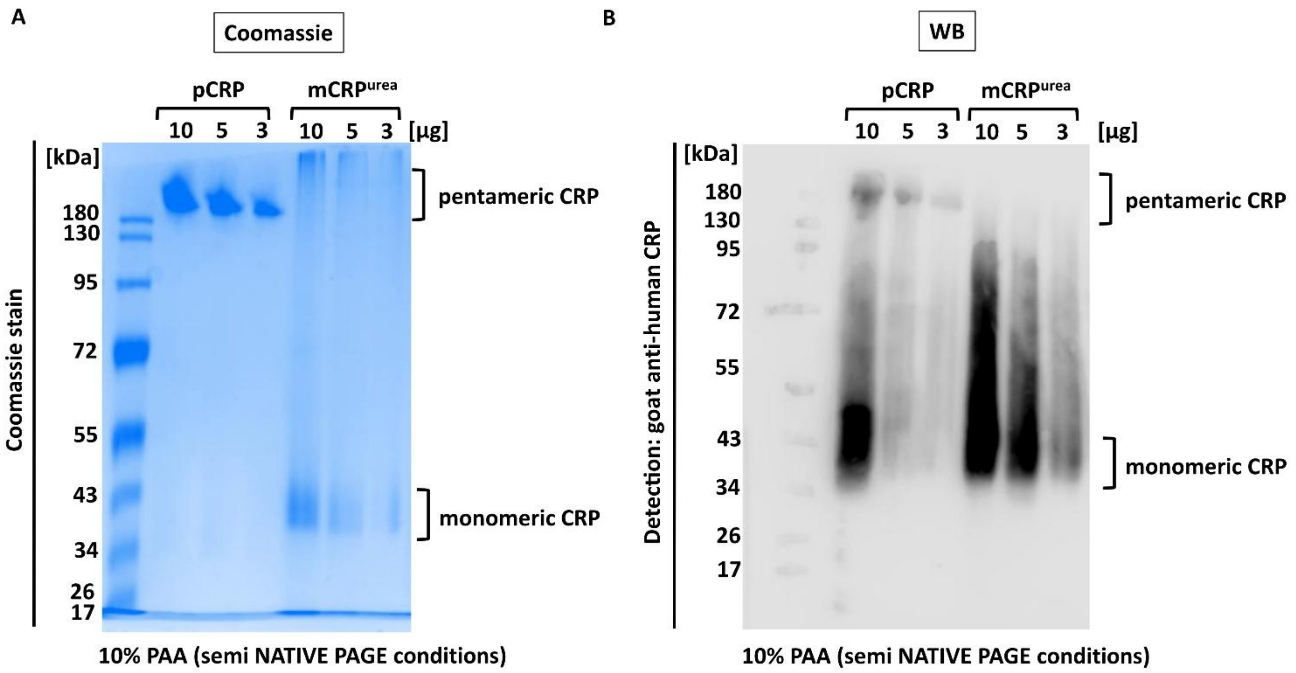


**Supplementary Figure 3:** Semi-native coomassie stain and western blot image probed for pCRP and mCRP conformation. In brief, samples were mixed with 15 µl of 1x SB (recipe, 1/20 of SDS, no DDT, no ß-ME) and pCRP or mCRP as indicated (10 µg, 5 µg or 3 µg). Samples were left without heating or boiling of the samples and loaded onto 10% PAA-Gel (all gel components and 1-Lämmli-Running buffer only with 1/20 of 20% SDS; final SDS Conc. 1%) and separated accordingly at 20 mA for ~ 1.5h. Gel was either directly stained by Coomassie-brilliant blue solution and destained with water (**A**) or transferred onto nitrocellulose membrane for western blot analysis. Here, membrane was stained with goat anti-human CRP followed by Donkey anti-goat IgG-POD and development by ECL substrate. Membrane exposure was developed using Licor Odyssey Imaging system) (**B**).
